# Automated multimodal imaging of *Caenorhabditis elegans* behavior in multi-well plates

**DOI:** 10.1101/2024.02.09.579675

**Authors:** Hongfei Ji, Dian Chen, Christopher Fang-Yen

**Affiliations:** Department of Biomedical Engineering, College of Engineering, The Ohio State University, Columbus, OH 43210, USA; Department of Bioengineering, School of Engineering and Applied Science, University of Pennsylvania, Philadelphia, PA 19104, USA

**Author notes:** **Corresponding author:** Christopher Fang-Yen.

**Keywords:** *C. elegans*, locomotion, egg-laying

## Abstract

Assays of behavior in model organisms play an important role in genetic screens, drug testing, and the elucidation of gene-behavior relationships. We have developed an automated, high-throughput imaging and analysis method for assaying behaviors of the nematode *C. elegans*. We use high-resolution optical imaging to longitudinally record the behaviors of 96 animals at a time in multi-well plates, and computer vision software to quantify the animals’ locomotor activity, behavioral states, and egg laying events. To demonstrate the capabilities of our system we used it to examine the role of serotonin in *C. elegans* behavior. We found that egg-laying events are preceded by a period of reduced locomotion, and that this decline in movement requires serotonin signaling. In addition, we identified novel roles of serotonin receptors SER-1 and SER-7 in regulating the effects of serotonin on egg laying across roaming, dwelling, and quiescent locomotor states. Our system will be useful for performing genetic or chemical screens for modulators of behavior.

## INTRODUCTION

*C. elegans* behavioral assays are widely used to study biological mechanisms in diverse areas such as aging (1, 2), sleep (3), development (4), and neurological disorders (5, 6). Assays of worm behavior have traditionally been done by manual assays and visual observations, which are labor intensive, subject to observer error, and low in throughput due to examination of a single animal at a time.

To address these limitations, researchers have developed systems for analyzing worm behaviors via video recording and software analysis (7, 8). These include methods for imaging worms on standard agar plates (9–15), in microfluidic devices (16–21), and in multi-well substrates (1, 22–26).

Our laboratory has been conducting behavioral screens for the effect of genetic mutations and/or pharmacological compounds on various aspects of *C. elegans* behavior, including locomotion, egg laying, and behavioral states such as quiescence, dwelling, and roaming (1, 15, 21, 23, 27, 28). Toward this end, we sought to develop an efficient method for screening a large number of conditions for multiple behaviors simultaneously.

Here, we report a method for multimodal behavioral recording and analysis of *C. elegans* using 96-well plates. We developed a computer vision software toolbox to automate analysis of several behavioral metrics including locomotion activity, behavioral states, locomotor frequency, and egg laying. This automated approach is capable of continuously recording behaviors from 96 individual animals simultaneously for several hours under controlled conditions. We demonstrate the capabilities of this system by analyzing the roles of serotonin signaling in locomotor and egg-laying behaviors. Our findings underscore the extensive capabilities of longitudinal imaging systems for large-scale phenotypic assays.

## METHODS

### C. elegans strains and maintenance

*C. elegans* strains were cultivated on *Escherichia coli* strain OP50 at 20 °C using standard conditions (29). The strains used in this work were obtained from the NIH *Caenorhabditis* Genetics Center. Mutant strains were the following: MT1082 *egl-1(n487)*, DA1814 *ser-1(ok345)*, DA2100 *ser-7(tm1325)*, DA2109 *ser-7(tm1325); ser-1(ok345)*. All experiments were performed with 1-day-old adult animals in a food-free environment. Worm populations were synchronized by timed egg lays according to standard methods (30).

### Multi-well plate assays

To study the effect of various chemicals on egg-laying and locomotion behaviors, we used 96-well microtiter assay plates (Corning 3795, round bottom), with each well containing the indicated number of worms immersed in 60 μL of NGM buffer dissolved with the compound of interest. The liquid NGM buffer consists of the same ingredients as solid NGM but without agar, peptone, or cholesterol (31).

### Image data acquisition and processing

The assay plate was placed on a platform within a custom-built imaging system. Dark-field illumination was generated by an LED light ring (outer diameter 12 inches; Sunpak) above the platform. A video was captured using a CMOS camera (DMK 33UX183, The Imaging Source) with a C-mount lens (Fujinon HF12.5SA-1, focal length 35 mm, f/2.8 or Nikon, focal length 55 mm, f/3.5 F-mount lens with a C-mount adapter) for 20 s at 5 frames per second using IC Capture software (The Imaging Source). A whole course of behavioral assay was performed by capturing a sequence of the 20 s videos, taken every 2 min for 5 h of recording time. The field of view of the system was set to display all 96 wells of the plate, which yields an 18 μm pixel resolution.

Video imaging data from the assay plate was analyzed using custom-written MATLAB and Python software. The behavioral outcomes from the analysis include locomotion activity, locomotor frequency, and egg-laying behavior. Locomotion activity is defined as the number of pixels whose intensity changes between subsequent image frames due to worms’ movements (1, 3). Locomotor frequency is calculated by our automated image analysis software capable of rapidly estimating rhythmic behaviors (32). Egg-laying behavior is quantified by the number of eggs released into the environment. Eggs are detected by a trained mask-regional convolutional neural network (Mask-RCNN) (33).

### Measuring locomotory behavior

We calculated locomotor activity using previously described methods (3, 27). Pairs of temporally adjacent video images were subtracted to generate difference images, which were then divided by the average pixel intensity, yielding normalized maps of pixel value intensity change. A Gaussian smoothing filter with a standard deviation of one pixel was applied to decrease image noise. A binary threshold was applied to the filtered image to determine the presence or absence of movement at each pixel location. The sum of all pixels exhibiting movement was calculated as the locomotor activity.

To calculate locomotor frequency we used our custom algorithm Imaginera (<u>imag</u>e <u>in</u>variants <u>e</u>nsemble for rhythmicity <u>a</u>nalysis) (32). For each video frame, our algorithm estimates the pixel area of a worm (*A*_*worm*_) in each well. When animals approached the edge of wells, they were occasionally occluded such that the animals could not be easily observed, decreasing *A*_*worm*_. To mitigate the influence of this occlusion, we censored any frames during which *A*_*worm*_ was smaller than a preset empirical threshold when calculating locomotor activity or frequency. For all the image data used in this study, the number of censored images within an individual videos was less than 0.2% on average and less than 5% at maximum.

### Measuring egg-laying behavior

To evaluate egg-laying behavior, we counted the eggs that have been released into each well in 20 s videos. Each video was sampled by 5 images evenly spaced across the 20 s interval, with adjacent images separated by 4 s. Through manual counting, the events where eggs were occluded by moving worms were counted to yield a frequency less than 0.3%.

Following previously described methods (34), we deployed a mask-regional convolutional neural network (Mask-RCNN) to detect and identify eggs in each image. The Mask-RCNN model was trained, evaluated, and deployed by PyTorch GPU (v2.0.0 + cu1.1.8) on a system equipped with a 13th Intel(R) Core(TM) i7-13700K processor and an NVIDIA GeForce RTX 4080 GPU.

For the model training, 1653 cropped single-well images (containing eggs and worms) from 18 videos in all 96 distinct well positions were manually annotated and saved as training datasets using a Python annotation package. We augmented the training set using operations including flipping, intensity adjustment, and Gaussian blurring, yielding a total of 6612 annotated images in the dataset. The model was trained for 25 epochs, during which the training set was randomly divided into training and testing sets with an 80/20 split ratio (5290 images for training, 1322 images for testing). We evaluated the Mask-RCNN model using characterization metrics, including precision (a measure of false positive rate), recall (a measure of false negative rate), and average precision (a measure of overall accuracy). Specifically, the RCNN model yielded 0.85 in precision, 0.81 in recall, and 0.75 in average precision; these metrics were similar to those of previous RCNN models used for *C. elegans* egg detection (35).

With the trained Mask-RCNN model, the egg number in an image was determined by *N*_*RCNN*_ = *A*_*RCNN*_/*a*_0_, where *A*_*CNN*_ represents the pixel number of egg objects inferred by the model, and *a*_0_ is the pixel number of a single egg, estimated by 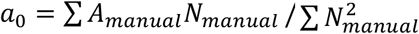. Here, ∑ *A* _*manual*_*N*_*manual*_ is the sum of the products of egg pixel number and egg number manually counted from individual images in the training set, and 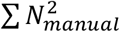 is the sum of squares of egg numbers from the same set.

### Mask R-CNN model characterization

We evaluated how well our model performed by calculating precision and recall as well as average precision (AP). Precision is the proportion of true positive predictions (correctly detected objects) among all positive predictions (correctly detected objects plus false positives). Recall is the proportion of true positive predictions among all actual positive instances (true positives plus false negatives). Average precision was calculated by computing the area under the Precision-Recall curve through the following steps: (a) For each confidence threshold, calculate precision and recall. (b) Interpolate the precision values by taking the maximum precision for any recall value greater than the current recall value. (c) Compute the area under the interpolated Precision-Recall curve using numerical integration.

To determine true positives (TP), false positives (FP), and false negatives (FN), we used the Intersection-over-Union (IoU) threshold, which measures the overlap of the ground truth bounding box and the prediction bounding box (35). An egg detection is considered a TP if its IoU with the corresponding ground-truth bounding box is greater than or equal to the IoU threshold. If the IoU is below the threshold, it is considered an FP. An FN is a ground-truth bounding box that does not have a corresponding detection. An IoU threshold of 0.3 and a confidence score threshold of 0.01 were used for our egg-detection model.

### Experimental design and statistical analyses

Differences in locomotion activity, active state fraction, locomotor frequency, and egg-laying rate (Figs. 3 I-L, Figs. 5 C-F) between various conditions were determined using the two-tailed Student’s *t*-test. Comparisons of serotonin resistance between different strains were performed using the two-way ANOVA test (Figs. 3 I and J). Differences in midpoint egg-laying time distributions (Fig. 3M) were assessed using a Wilcoxon rank sum test.

## RESULTS

### System design for high-content imaging of multi-well assay platform

Our imaging platform contains a 20-megapixel CMOS camera, a C-mount lens, a 30 cm diameter LED ring illuminator, and a plate holder (Fig. 1A). One or more worms in liquid NGM media are prepared in each well of a 96-well plate and the plate is imaged under dark field illumination (Figs. 1 A and B). The platform components are positioned to visualize worms and embryos in each well while minimizing stray light from well edges (Fig. 1B).

**Figure 1.**
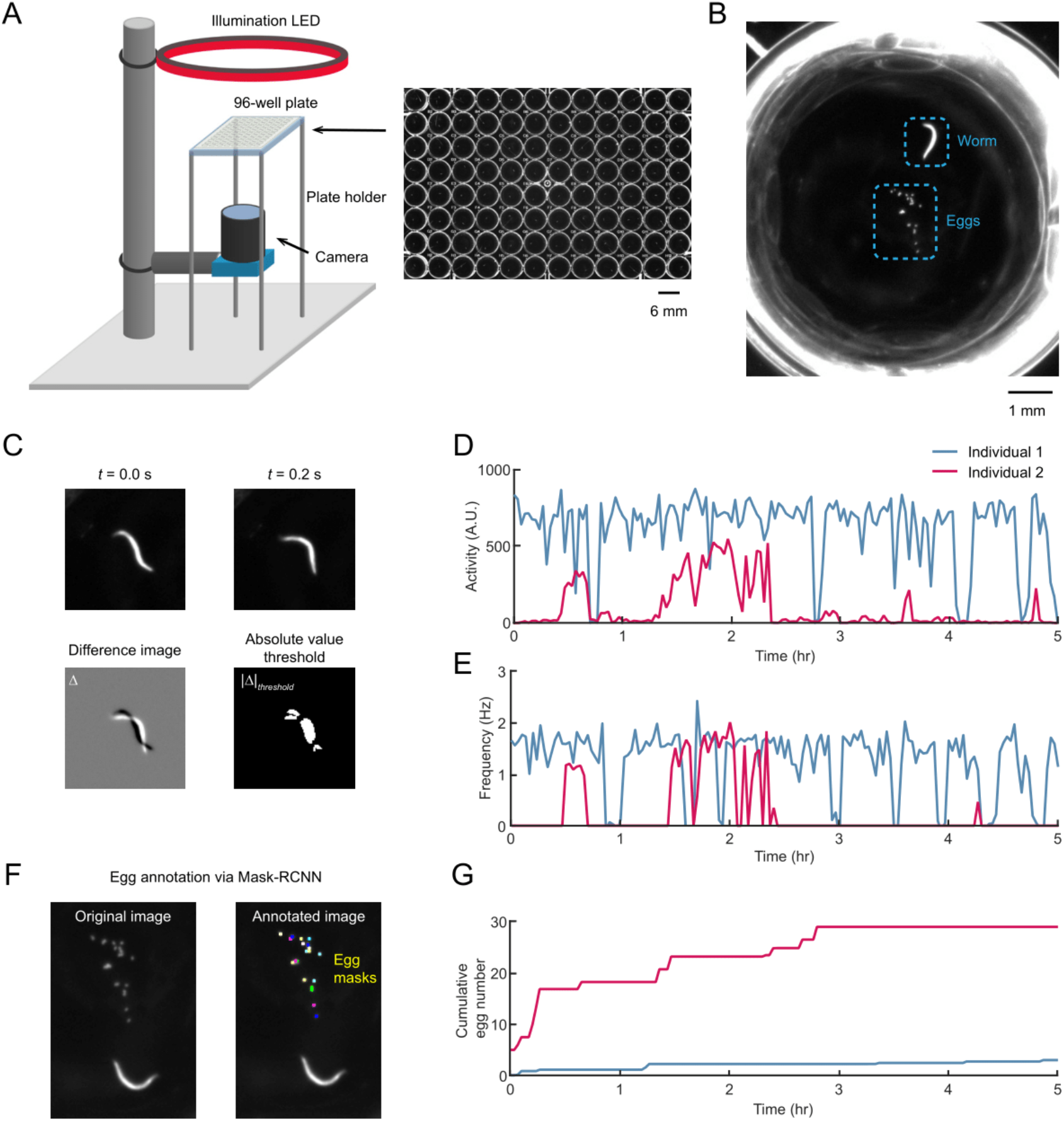
System design and behavioral quantification. (A) Schematic of the imaging system and image of a 96-well assay plate containing one worm per well. (B) Detail of a well containing a single 1-day adult and multiple eggs. (C) Activity is calculated as the sum of the number of pixels whose pixel value intensities change above a threshold. The change in the pixel value intensity is calculated by the pixel-by-pixel difference of temporally adjacent images. (D) Activity over time of two individual wild-type animals. (E) Locomotor frequency over time of two wild-type animals. (F) Egg detection via Mask R-CNN from the original video frame (*Left*). The egg objects are annotated with masks inferred by the model (*Right*). (G) Number of eggs laid for two wild-type animals.

### Computer vision automates multimodal behavioral measurements

We developed image analysis software for quantifying animal locomotory and egg-laying behaviors within each well. Using a combination of computer vision algorithms and convolutional neural network (CNN), the software analyzes images to measure behavioral parameters, including locomotion activity, locomotor frequency, and egg-laying rate (see *Methods*).

Locomotion activity is computed through pixel-by-pixel subtraction of consecutive video frames, with the activity defined by the number of pixels exhibiting intensity changes between subsequent frames (Figs. 1 C and D) (3). To measure locomotor frequency, we used an algorithm based on analysis of image invariants (Fig. 1E) (32). The image analysis software also employs a Mask-Region Convolutional Neural Network (Mask-RCNN) to automatically annotate and enumerate worm embryos in each well (Figs. 1 F and G) (33, 34).

### Evaluation of automated measurements

We tested the accuracy of our software in analyzing the locomotor frequency and egg-laying events by comparing the software’s measurements with those obtained through human observation.

The automated frequency results demonstrated good agreement with manual data with an overall root mean squared error of 0.097 Hz and an R-squared value of 0.964 (32). We show that compared to previous established computer-vision approaches (e.g., segmentation-based method), our technique is more robust to low image quality and background noise (32). We found that locomotion activity is also highly correlated with locomotor frequency by comparing it with the ground-truth data (Fig. 2A).

**Figure 2.**
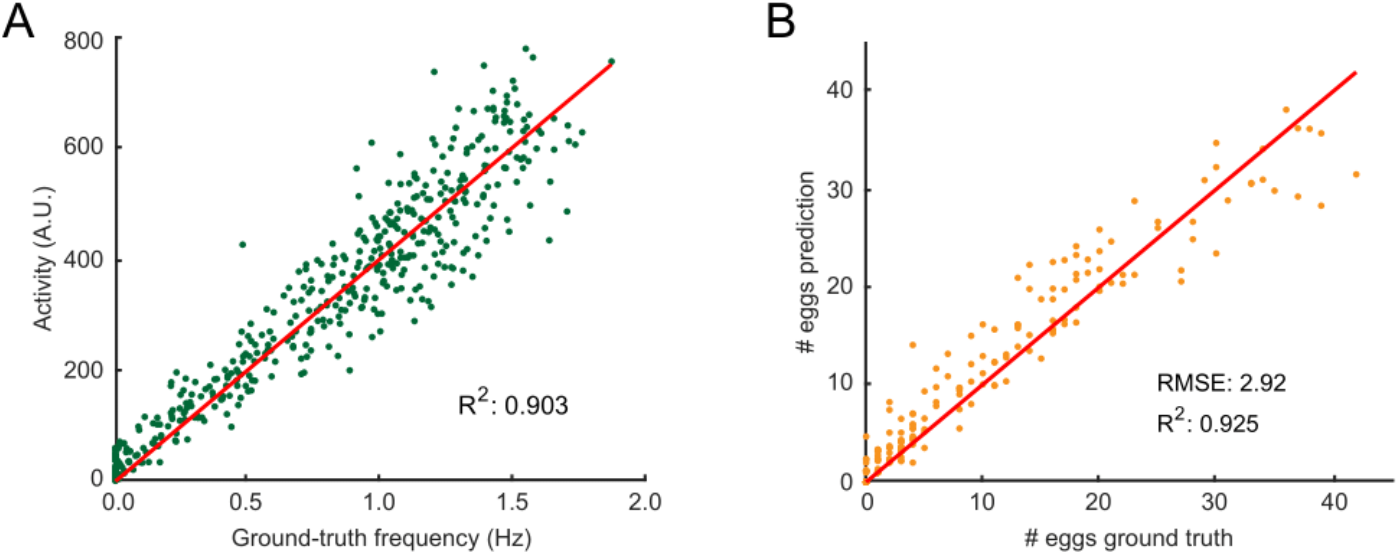
Evaluation of automated behavioral measurements. (A) Locomotion activity compared with the frequencies calculated by manual observation of recorded videos. The line indicates a linear fit with zero offset. (B) Egg numbers inferred by the Mask-RCNN model compared with those counted manually. The line shows equality between predicted and ground truth numbers.

**Figure 3.**
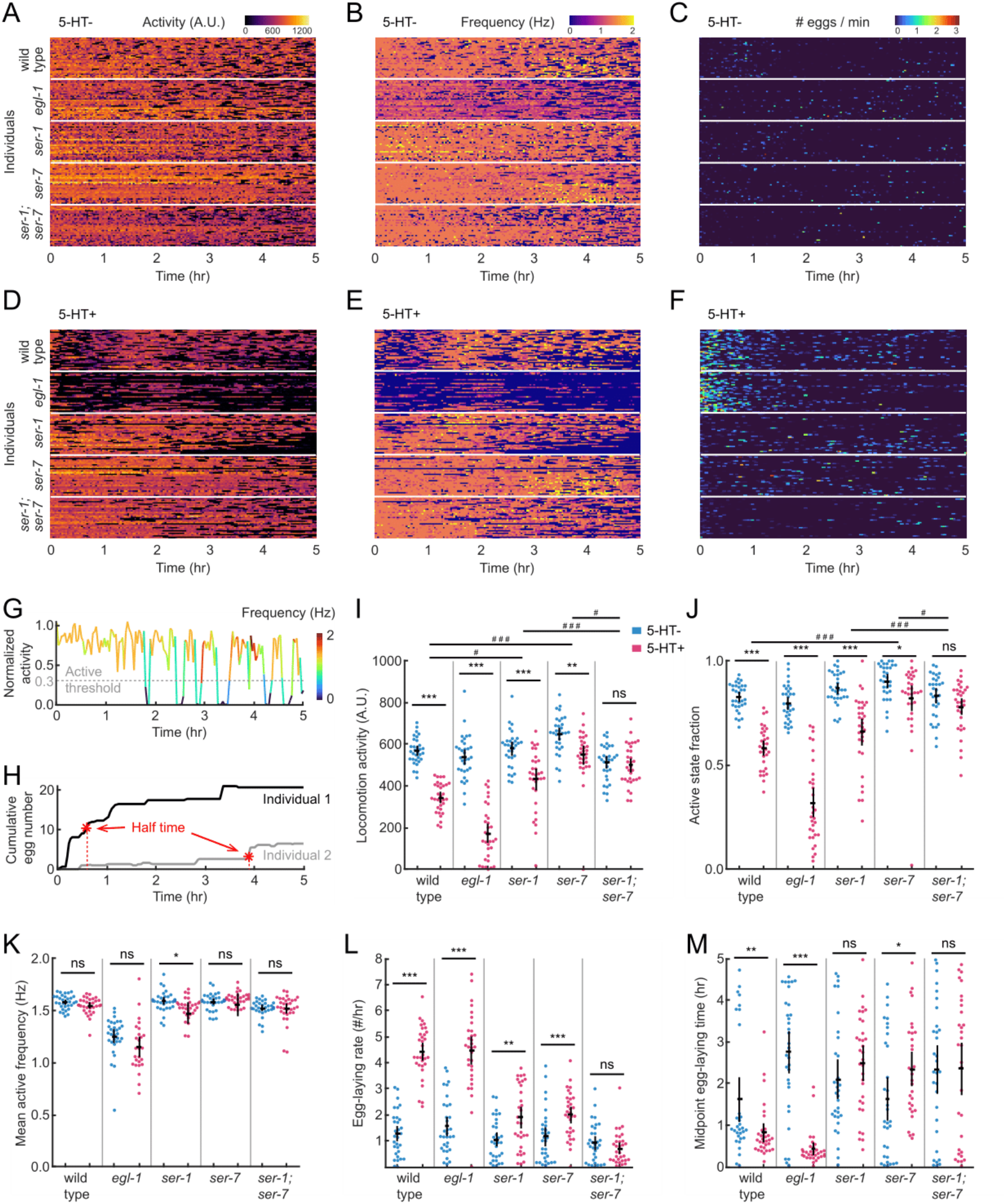
Automated large-scale tracking of locomotory and egg-laying behaviors. (A-C) Heat maps of activity (A), locomotor frequency (B), and egg-laying rate (C) for wild-type and mutant individuals (n = 32 for each strain group). Each row in the heatmap shows the behavioral dynamics of a single animal. (D-F) Same as (A-C), but for animals in an exogenous serotonin environment. (G) Normalized activity over time for an individual. The data is normalized by the mean value of the 95^th^ to 100^th^ percentile of all activity values observed during the experiment. Color represents the locomotor frequency of the individual. Gray dashed line indicates the threshold value of normalized activity above which locomotion bouts are labeled as active states. (H) Cumulative number of eggs released by two individuals. Red asterisk indicates midpoint egg-laying time (*t*_50_), the time taken to reach one-half of the total egg number accumulated at t = 5 h. (I) Locomotion activity of wild-type and mutant individuals. (J) Fraction of time during active state of wild-type and mutant individuals. (K) Mean locomotor frequency during active state of wild-type and mutant individuals. (L) Egg-laying rate of wild-type and mutant animals. (M) Midpoint egg-laying time (*t*_50_) of wild-type and mutant individuals. In (I-M), each point represents an animal, red and blue indicate groups in the respective serotonin conditions, black bars represent population mean ± SEM. *p < 0.05, **p < 0.01, ***p < 0.001, Student’s *t*-test. In (I-J), several indicated dosed/undosed pairs of strains were compared to further investigate the interaction between the genetic and chemical factors. #p < 0.05, ###p < 0.001, two-way ANOVA.

To assess the accuracy of the egg-detector RCNN model, we applied it to identify and count eggs across all 96 wells of the plate. The collected well images contained 1-day-old adult wild-type worms alongside the eggs they laid, captured at 2 and 5 hours after animals were transferred into the assay plate. We found that the egg counts inferred by the model were in good agreement with those counted manually (Fig. 2B).

Together, the automated methods of our system accurately measure locomotion activity, locomotor frequency, and embryo numbers through longitudinal imaging.

### Automated tracking of locomotion, behavioral states, and egg laying

*C. elegans* locomotory and egg-laying behaviors are modulated by serotonergic, dopaminergic, and peptidergic signaling (36–38). The *C. elegans* nervous system uses serotonin (5-HT) to modulate both its locomotory and egg-laying behaviors, and external serotonin has been found to inhibit locomotion and stimulate egg-laying activity (23, 39–41).

To evaluate our system’s reliability and capability for large-scale assays, we tracked the locomotion and egg-laying behaviors of wild-type and mutant *C. elegans* individuals under controlled serotonin conditions (see *Methods* and Figs. 3 A-F). Animals were individually housed in wells of an assay plate filled with (food-free) liquid NGM and defined serotonin concentrations (5-HT-: 0 mM; 5-HT+: 6.5 mM). In each experiment, we monitored up to 96 animals for 5 h, which yielded 151 video clips with 5 fps and 20 s duration.

Both wild-type and mutant animals under serotonin-free conditions showed relatively active locomotion (Figs. 3 A and B) and low egg-laying rates (Fig. 3C). In contrast, the administration of exogenous serotonin induced diverse behavioral dynamics across the tested strains (Figs. 3 D-F).

We defined parameters for quantifying each animal’s locomotory states (42). We calculated the mean of each animal’s higher 5% activity values across the experiment and used this value to define a normalized activity. This data allowed us to identify sedentary and active bouts of locomotion using an empirical threshold (Fig. 3G) (23). We focused on assessing the average locomotion activity, the fraction of time spent in the active state, and the average locomotor frequency during the active state.

Wild-type and mutant animal populations exhibited diverse locomotion activity levels and active locomotory states across genotypes and different levels of exogenous serotonin (Figs. 3 I and J). Both wild-type and mutant animals generally exhibited high activity levels and spent the majority of time (over 78%) in the active state under the serotonin-free condition (Figs. 3 I and J). With the addition of exogenous serotonin, activity levels and active state fractions decreased in wild-type animals (reduced by 39% and 29% on average, respectively) and in *egl-1* mutants (which lack HSN serotonergic neurons that innervate egg-laying muscles; reduced by 68% and 60%, respectively) (Figs. 3 I and J). In contrast to the wild type, *ser-1* and *ser-7* mutants (deficient in G protein-coupled metabotropic receptors SER-1 and SER-7) both exhibited resistance to the effects of serotonin on locomotion activity (reduced by 25% and 15% for the respective mutants) and active state fractions (reduced by 24% and 9% for the respective mutants), while *ser-1;ser-7* double mutants showed enhanced resistance to exogenous serotonin (2% reduction in locomotion activity, 6% reduction in active state fraction) (Figs. 3 I and J). This result suggests that the SER-1 and SER-7 receptors play a role in mediating inhibitory effects of external serotonin on locomotor behavior.

We calculated the average locomotor frequency of each animal during its active state. The average active locomotor frequency of wild-type and all tested mutant strains showed little or no change in response to exogenous serotonin (Fig. 3K). Therefore, unlike the active state fraction, the active locomotor frequency remained largely constant, independent of exogenous serotonin. This experiment suggests that external serotonin on locomotion primarily affects initiation of activity rather than alteration of an established locomotor rhythm.

To characterize the egg-laying dynamics of each animal, we calculated the average egg-laying rate and the midpoint egg-laying time *t*_50_, the time taken to reach half of the total number of eggs laid during the experiment (Fig. 3H). Under serotonin-free conditions, wild-type animals averaged an egg-laying rate of 1.2 eggs/h (Fig. 3L). The population’s *t*_50_ values exhibited a biphasic distribution, with the majority displaying early midpoint egg-laying times (Fig. 3M). Compared to the wild type, the mutant animals tested had similar egg-laying rates but displayed different distributions in *t*_50_ (Figs. 3 L and M). In the presence of exogenous serotonin, wild-type and *egl-1* mutant animals showed significant increases in egg-laying rates and decreases in *t*_50_ values (Figs. 3 L and M). The *ser-1* and *ser-7* mutants had a moderate increase in egg-laying rates in response to exogenous serotonin, while *ser-1;ser-7* double mutants maintained low egg-laying rates (Fig. 3L). The values of *t*_50_ in the wild-type and *egl-1* mutant animals were suppressed with the addition of exogenous serotonin, which was not seen in the *ser-1, ser-7*, and *ser-1;ser-7* double mutants (Fig. 3M). This result indicates that SER-1 and SER-7 receptors may be necessary for normal serotonin-mediated egg laying involving both egg-laying rates and temporal patterns.

In summary, our behavioral findings qualitatively agree with previously reported results for each strain, encompassing behavioral parameters such as locomotory state, locomotor frequency, and egg-laying rate, as well as the effects of exogenous serotonin (4, 22, 27, 39, 41, 43–45). These experiments demonstrate our system’s reliability, scalability, and efficiency in concurrently and longitudinally assessing multi-metric behavioral dynamics.

### Serotonin is required for preparatory movement decline during egg-laying

One advantage of our system is that it is capable of capturing locomotor and egg-laying behaviors simultaneously. We examined the temporal correlation between locomotion and egg-laying by extracting the locomotion pattern before and after each egg-laying event throughout the recording period (Figs. 4 A and B). The average locomotor activity of wild-type animals was found to decrease approximately 10 min prior to egg-laying events, with the minimum activity reached roughly at the time the egg(s) were laid and subsequently recovering to a baseline value within about 10 min (Fig. 4C). Observation of the video clips revealed that the activity of individual animals during egg-laying events was consistently lower than during adjacent periods during which no eggs were laid (individual traces in Fig. 4C). Notably, nearly every egg-laying event was preceded by a decline in activity, although the magnitude and timing of the maximal activity drop varied substantially both within individual animals and among different animals (Fig. 4C).

**Figure 4.**
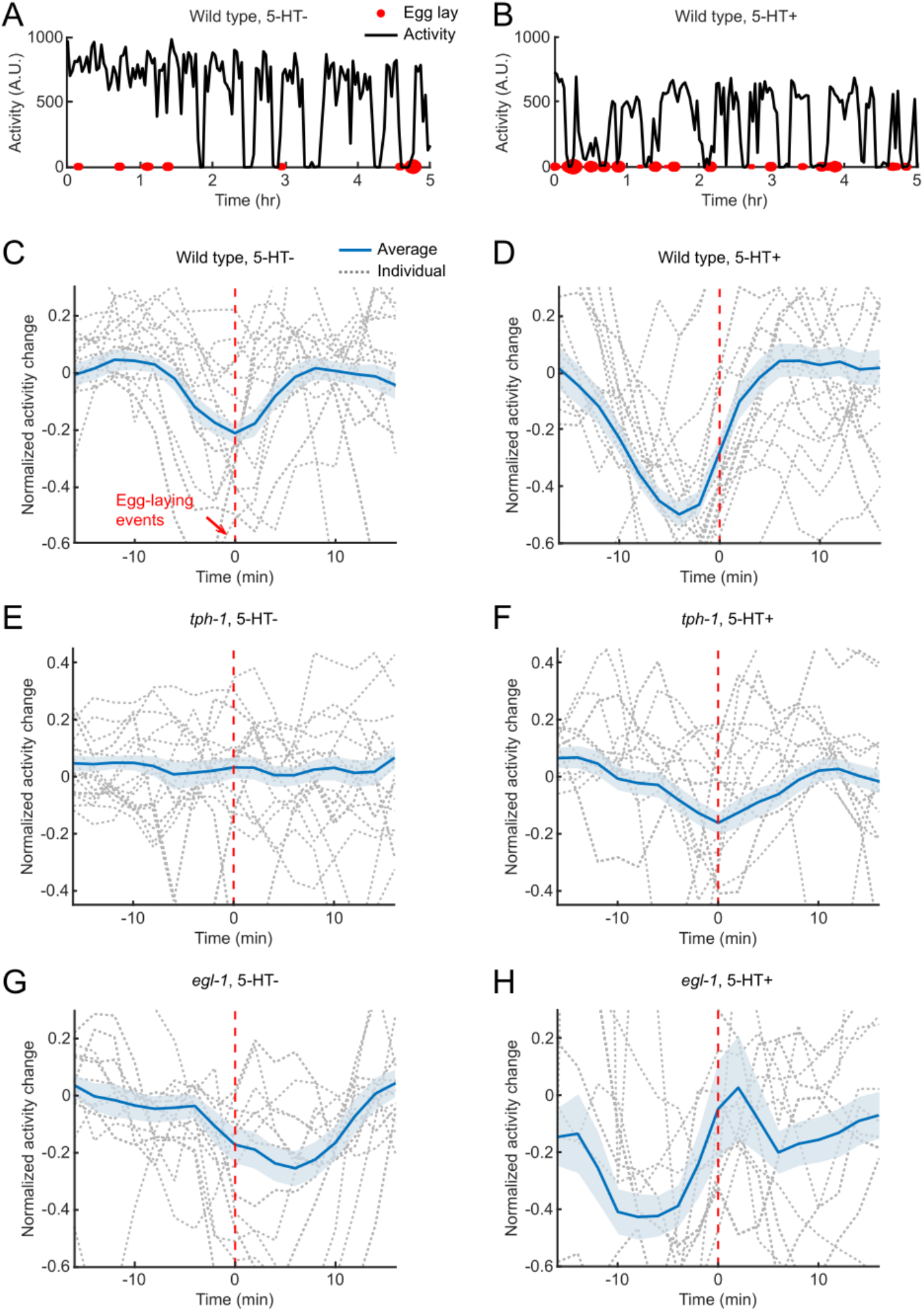
Serotonin is required for preparatory movement decline during egg laying. (A and B) Locomotion activity and egg-laying dynamics for two individual wild-type animals in either the absence (A) or presence (B) of exogenous serotonin (B). Black curves represent activity trajectories. Red dots indicate egg-laying events scored by automated method, and dot size is used to visualize the egg-laying rate. (C and D) Average normalized activity change around egg-laying events (red dashed line, aligned at *t* = 0) of wild-type animals (n = 32) in either the absence (C) or presence (D) of exogenous serotonin. Blue line and shaded area indicate population mean ± SEM. Gray dashed curves represent 20 randomly selected individual trajectories for each group. (E and F) Same as (C and D), but for *tph-1* mutant animals (n = 32). (G and H) Same as (C and D), but for *egl-1* mutant animals (n = 32).

**Figure 5.**
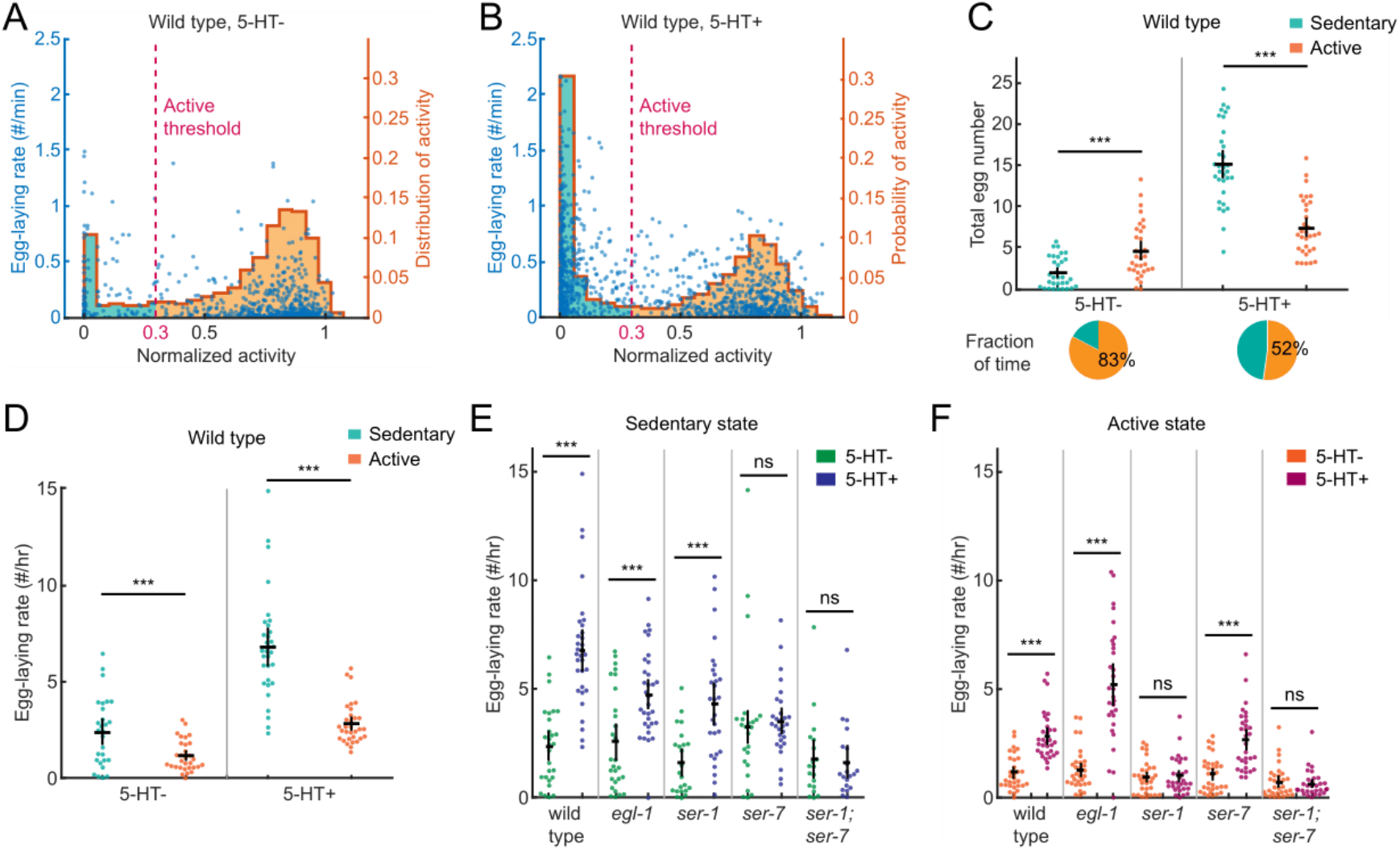
Receptors SER-1 and SER-7 regulate serotonin effects on reproduction in separate locomotory states. (A and B) Histogram of normalized activity (orange stairstep curve, right-side y-axis) and scatter plot showing egg-laying rate versus normalized activity value (blue dot, left-side y-axis) for wild-type animals in either the absence (A) or presence (B) of exogenous serotonin. In histograms, the height of the stairstep denotes the relative probability of each of 20 bins of normalized activity, red dashed line indicates the threshold value of normalized activity above which locomotion is flagged as active state, green and orange shaded areas represent proportions of sedentary and active states. In scatter plots, each point represents a 20 s bout of an individual animal (n = 4832 bouts from 32 animals for each condition). (C) (*Upper*) Total number of eggs released by wild-type animals during sedentary or active states in either the absence or presence of exogenous serotonin. (*Lower*) Fraction of time of sedentary versus active states in respective serotonin conditions. (D) Egg-laying rate of wild-type animals during sedentary or active states in either the absence or presence of exogenous serotonin. (E and F) Egg-laying rate of wild-type and mutant individuals during sedentary (E) or active (F) state in either the absence or presence of exogenous serotonin. In (C-F), each point represents an animal, and black bars represent population mean ± SEM. ***p < 0.001.

Since serotonin functions in circuits that control *C. elegans* locomotion and egg laying (4, 39, 41, 42, 46), we examined the effects of serotonin and the neurons regulating egg-laying behavior on the control of locomotion. We asked how exogenous serotonin might modulate the observed correlation between locomotion and egg-laying. In the presence of exogenous serotonin, the decline in movement around egg-laying events was more pronounced. Furthermore, the timing of the activity decline and subsequent recovery occurred approximately 4 min earlier than without exogenous serotonin (Fig. 4D).

Next, we explored the effects of endogenous serotonin signaling on the coordination between egg-laying and locomotion. We analyzed mutant animals deficient in serotonin due to a mutation in the biosynthetic enzyme tryptophan hydroxylase (TPH-1). While the overall locomotion of *tph-1* mutants appeared normal (14), these mutants did not exhibit a decline in movement around egg-laying events (Fig. 4E). The introduction of exogenous serotonin restored this coordination deficit (Fig. 4F). Together, these results show that serotonin signaling is required for the observed decline in movement during egg-laying events.

We further investigated the effects of neurons that regulate egg-laying behavior on the coordination with locomotion, focusing on the role of HSNs, a pair of serotonergic neurons crucial for a normal egg-laying process. We observed the behavior of *egl-1* mutants, which specifically lack HSNs. While *egl-1* mutants did exhibit a decline in movement around egg-laying events, the timing of their minimum activity was delayed, occurring consistently after egg-laying (Fig. 4G). Administering exogenous serotonin to *egl-1* mutants altered the timing of the movement decline (Fig. 4H). These results suggest that HSNs may be bypassed with the administration of exogenous serotonin for restoring the coordination between egg laying and locomotion.

### The metabotropic serotonin receptors SER-1 and SER-7 regulate serotonin effects on reproduction in separate locomotory states

Our analyses of *C. elegans* locomotion activity and behavioral states showed that animals occasionally alternated between sedentary and active states (Figs. 3 A, D, and G). This alternation between states was influenced by exogenous serotonin and seemed to happen at a frequency similar to that of egg-laying events (Figs. 4 A and B). We asked whether there is a correlation between *C. elegans* egg-laying behavior and behavioral states and whether one is influenced or modulated by the other.

We discerned sedentary and active periods from the locomotion activity trajectories of individual animals and registered these periods with their respective egg-laying rates (Figs. 5 A and B). We noted that in serotonin-free environments, wild-type animals spent the majority of their time in the active state (Fig. 5A) and laid more eggs during these active periods (Fig. 5C). Conversely, in environments with exogenous serotonin, animals spent nearly equal amounts of time on sedentary and active states (Fig. 5B), with more eggs being laid during the sedentary states (Fig. 5C).

To compare the egg-laying activity between different locomotory states, we calculated the egg laying rate by dividing the total egg count during each state with the corresponding state duration. The data revealed that, irrespective of exogenous serotonin, wild-type animals consistently exhibited higher egg-laying rates during sedentary locomotor state (Fig. 5D).

The observed dual role of serotonin in regulating both locomotory states and egg-laying behavior (as shown in Figs. 5 A and B) prompted a closer examination of serotonin signaling in modulating the observed dependence of egg-laying behavior on locomotory states (Figs. 4C and 5D). We revisited the behavioral recordings of HSN-defective mutants (*egl-1*) and serotonin receptor mutants (*ser-1, ser-7*, and *ser-1;ser-7*), with a particular focus on the effects of exogenous serotonin on the egg-laying rate during distinct locomotory states. By registering the egg-laying rate over time for each mutant individual and correlating it with the respective locomotory states, we observed that both wild-type and *egl-1* mutant animals experienced a significant increase in egg-laying rate during both sedentary and active states when exposed to exogenous serotonin (Figs. 5 E and F). Interestingly, the *ser-1* and *ser-7* mutants resisted the effects of exogenous serotonin on their egg-laying rates, but each during one of the two states (Figs. 5 E and F). Meanwhile, the *ser-1;ser-7* double mutants resisted the egg-laying effects of exogenous serotonin during both states (Figs. 5 E and F).

In summary, our findings suggest that *C. elegans* preferentially lay eggs during the sedentary state, with serotonin receptors SER-1 and SER-7 distributing the effects of serotonin on egg-laying behavior across distinct locomotory states.

## DISCUSSION

Our automated image recording and analysis system, set within the conventional 96-well plates, yielded substantial benefits in throughput, preservation of animal individuality, multi-metric behavioral output, and manageable and controlled external conditions. Our system facilitates the individual evaluation of parameters involving locomotion activity, behavioral states, and egg-laying behavior. This allows for the identification of diverse locomotion and reproduction characteristics across individuals and populations.

We obtained large-scale longitudinal behavioral recordings of individual animals through high-resolution imaging. The integrated computer-vision software toolkit automates quantifying various behaviors, leveraging accurate assessments of animal motility, locomotor rhythmicity, and egg laying. Applications enabled by our method included pharmacological screens (47), genetic screens related to diverse neural functions (48), and studies of complex behavioral patterns and states (49).

We observed that the algorithm for egg counting tends to slightly underestimate at higher egg numbers (Fig. 2B). We speculate that this may be due to the tendency for *C. elegans* egg laying to occur in short bursts, forming clumps of multiple eggs (50). In such a clump, eggs are likely to partially occlude each other, leading to an underestimate of egg numbers. To address this issue in the future, we may use an image dataset with a wider range of egg numbers to train a neural network model so that it detects individual eggs with higher accuracy in both sparse and crowded scenarios.

We observed that wild-type animals exhibit a decline in movement within a few minutes of egg-laying events. This finding diverges from those of previous analyses in which locomotion was observed to increase around egg-laying events (23, 44). This discrepancy may stem from variations in experimental food conditions; while our studies monitored animals in food-free environments, prior experiments involved animals on food, which potentially influenced their basal behavioral and physiological states (51). Future experiments could explore how different levels of food availability influence the coordination and adaptations of these behaviors.

Our automated imaging and analysis pipeline is scalable, platform-independent, and built from off-the-shelf parts. It is compatible with other standard or customized multi-array setups used for large-scale behavioral assays (1). Its modular analysis algorithm and customizable hardware configuration should allow for the integration of complex image-analysis routines specific to various screening tasks. For example, our system can be adapted for optogenetics-based screens by adding a high-intensity light source (1). The system can be suitable to taxis-based behavioral assays by using a customized plate with elongated lanes and controlled stimulus sources (e.g., chemical cues for chemotaxis) (26). Because of the system’s moderate dimension, it can fit in an intermediate sized thermal incubator and can potentially be used for temperature-dependent behavioral analyses. It also presents a versatile tool adaptable to studying not just various sizes of *C. elegans* but potentially also other small organisms like *Drosophila melanogaster* and its larvae.

One potential concern in any multi-well imaging setup is uneven variability of conditions across the plate, i.e. positional effects (52). These include nonuniform temperature, nonuniform dehydration, variability due to liquid handling procedures, temporal effects due to different loading times, nonuniform illumination, and the perspective effect, which causes wells near the edge of the field of view to appear slightly tilted compared to wells near the center. Common strategies for mitigating positional effects include avoiding using outer wells, complete or block-wise randomization of plate layout, and proper equipment calibration (52) The perspective effect can be reduced by increasing the distance between the camera and plate and using a longer focal length lens such that the plate continues to approximately match the size of the field of view; we have conducted experiments with lens focal lengths ranging from 35 mm to 75 mm. Another potential approach to reducing the perspective effect is to use a telecentric lens, which has a magnification independent of an object’s distance from the lens.

Our system is built from off-the-shelf components. A full parts list with installation instructions, analysis software, and test data are available at https://github.com/cfangyen/multiwell.

Our multi-array and multi-metric behavioral tracking automation will facilitate rapid execution of large, complex compound or genetic screens, making it possible to detect subtle phenotypes that are usually difficult to identify with single-metric behavioral tracking prevalent in existing systems. By assessing both locomotory and egg-laying activities, for example, one can screen for mutants that have altered coordination between the two behaviors or that exhibit subtle behavioral changes modulated by one or more signaling pathways.

## DATA AVAILABILITY

Strains are available upon request. The software and experimental sample data are available at https://github.com/cfangyen/multiwell.

## ACKNOWLEDGMENTS

We thank Niels Ringstad, Yen-Chih Chen, Mohammad Seyedsayamdost, Andrew Ruba, and Miriam Goodman for providing reagents and helpful advice. Some strains were provided by the *C. elegans* Genetic Center, funded by the NIH Office of Research Infrastructure Programs (P40 OD010440).

